# Evaluating the heterogeneous effect of extended incubation to blastocyst transfer on the implantation outcome via causal inference

**DOI:** 10.1101/2021.11.02.466894

**Authors:** Yoav Kan-Tor, Naama Srebnik, Matan Gavish, Uri Shalit, Amnon Buxboim

## Abstract

In IVF treatments, extended culture to single blastocyst-transfer is the recommended protocol over cleavage-stage transfer. However, evidence-based criteria for assessing the heterogeneous implications on implantation outcome are lacking. To estimate the causal effect of blastocyst-transfer on implantation outcome, we assembled a multicenter dataset of embryo time-lapse imaging. The data includes a natural source of randomness and has a strong claim for satisfying the assumptions needed for valid causal inference. By fitting a causal forest model, we assessed the ‘*Transfer Lift*’, which quantifies the probability difference in embryo implantation if transferred as a blastocyst versus cleavage-stage. Blastocyst transfer increased the average implantation rate, however we revealed a subpopulation of negative *Transfer Lift* embryos whose implantation potential is predicted to increase via cleavage-stage transfer. We provide day-of-transfer decision-support tools that are retrospectively estimated to improve implantation rate by 32%, thus demonstrating the efficacy of embryo-level causal inference in reproductive medicine.

**One Sentence Summary:** A causal inference model predicts the heterogeneous effect of prolonged incubation to blastocyst transfer on embryo implantation, thus providing means for optimizing pregnancy rates in IVF treatments.

## INTRODUCTION

During the past two decades, live-birth rates of *in vitro fertilization* (IVF) treatments have steadily declined worldwide,(*1*) concomitantly with the increase in maternal age.(*2*) The improvement in culture conditions supports extended embryo culture with single blastocyst transfer as an alternative to cleavage stage transfer,(*3*) thus decreasing the medical risks that are associated with multiple pregnancies.(*4-6*) However, inconsistent reports indicate that the causal effect of blastocyst transfer on embryo developmental competence remains unclear.(*7*) Randomized controlled trials (RCTs) suggest that blastocyst transfers can moderately improve live-birth rates only of fresh cycles but not the cumulative pregnancy rate per oocyte retrieval.(*8*) Despite being the method of preference for inferring causal relationships, RCTs are limited by methodological and ethical considerations, thereby hindering sample size, unbiasedness towards their target embryos, and representativeness of the true distribution of embryos.(*9*) Alternatively, large retrospective datasets of clinically-labeled video recordings of preimplantation embryo development are becoming increasingly available, owing to the utilization of time-lapse incubation systems in IVF clinics worldwide.(*10, 11*) Under certain special conditions, such datasets can generate compelling causal evidence that complements the evidence of RCTs.(*12*) Below we present a methodological and computational framework for evaluating the heterogeneous causal effect of extended incubation to blastocyst transfer on the implantation potential of embryos using observational data.

## RESULTS

### Assembling a retrospective dataset of embryo transfers and implantation outcome that provides a natural source of randomness and supports causal inference

To measure the effect of blastocyst-versus-cleavage stage transfer on implantation outcome while accounting for possible confounding factors, we assembled a retrospective multicenter dataset of clinically-labeled video files of preimplantation embryo development as reported previously.(*13*) The dataset includes 3,433 IVF cycles that were conducted during a five-year period. We focus on fresh transfer cycles of cleavage-stage embryos three days after fertilization (Day-3) and blastocysts five days after fertilization (Day-5), as illustrated in figure 1A. The embryos in the dataset were obtained from four independent large IVF clinics distributed across the country, which operate only on weekdays (Sunday-to-Friday). Since fertilization is performed on the same day as ovum pick up (OPU), Day-5 and Day-3 transfers become almost excluded for patients that underwent OPU on Mondays and Wednesdays, respectively (Fig. 1B). In turn, the day-of-OPU is determined according to follicular growth criteria, which lack a causal relationship with the potential implantation outcome of the transferred embryos.(*14*) Hence, day-of-OPU decision making provides a *natural source of randomness* with respect to the potential day-of-transfer outcome in our dataset. Importantly, infertility treatments are fully subsidized for the first and second children by the Israeli Ministry of Health. This policy contributes to a better *representation* of all socioeconomic and ethnical sectors of relevant age in our dataset.

**Figure 1.**
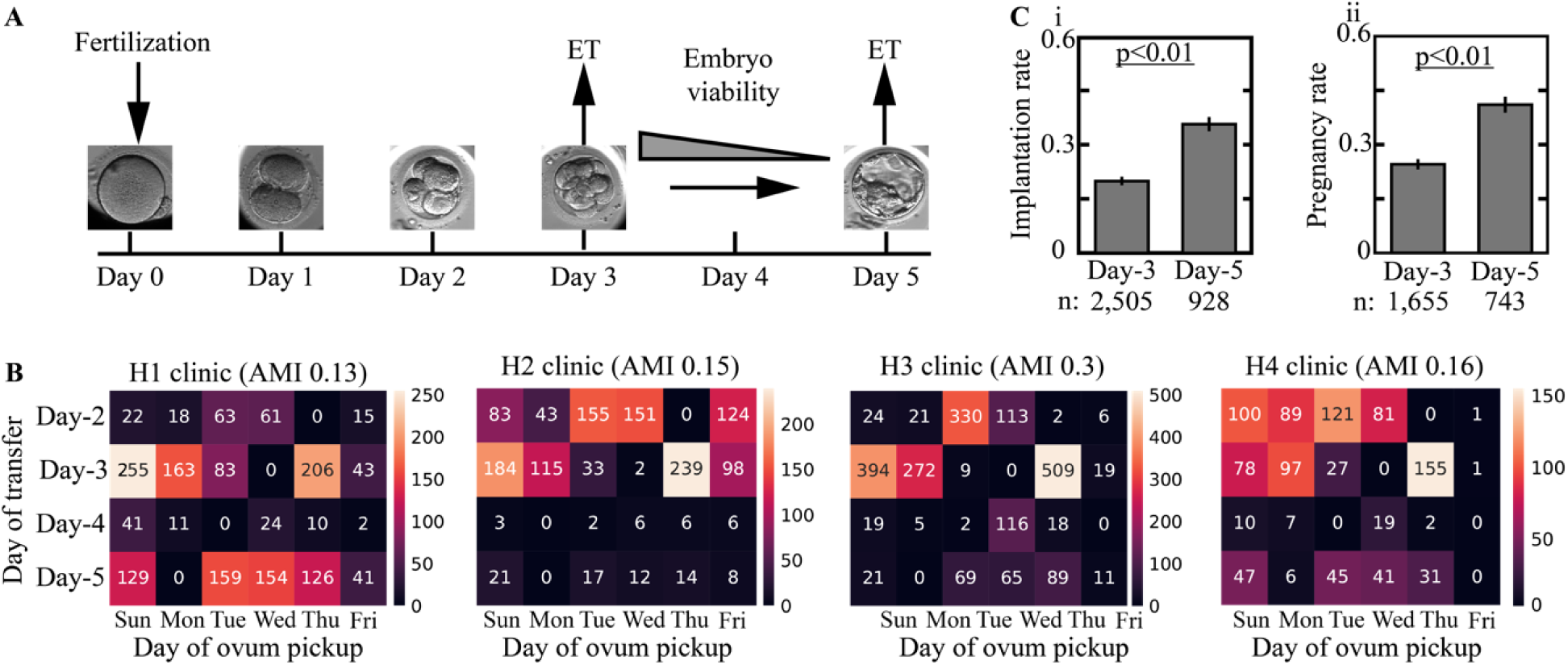
Delineating the causal model of cleavage-stage versus blastocyst transfers with respect to embryo implantation outcome in IVF-ET treatments. **A**, Embryos are transferred to the uterus either at cleavage stage (typically Day-3) or as blastocysts (typically Day-5). **B**, Day-of-transfer distributions are provided for all days-of-OP Sunday through Friday. Since all four clinics are closed on Saturdays, Day-3 and Day-5 transfers are excluded from OP’s that were performed on Mondays and Wednesdays, respectively. The statistical dependence between DoT and Day-of-OP distributions is quantified using the adjusted mutual information (AMI), which is indicative of a weak relationship. **C**, For fresh embryo transfers, Day-5 blastocyst transfers account for higher (i) implantation rate and (ii) pregnancy rate.

Our dataset indicates that the average implantation (Fig. 1C-i) and pregnancy (Fig. 1C-ii) rates of blastocyst transfers are 70% higher than cleavage-stage transfers. These differences can be attributed to de-selection of embryos that had exhibited normal cleavage on Day-3 from fertilization but became developmentally arrested and failed to blastulate on Day-5.(*7*) However, these differences cannot be taken to be causal due to potentially confounding factors in the treatment assignment process.(*15*) In order to estimate the causal effect of extended incubation to blastocyst transfer on implantation outcome from observational data, we will employ methods based on the idea of covariate adjustment (standardization).(*12*) As is well-established in the causal inference literature, for covariate adjustment to yield valid causal estimates, we must examine whether the data-gathering process satisfies the *ignorability* (unconfoundedness) and *overlap* assumptions, as well as the stable unit treatment value assumption (*SUTVA*).(*12, 16*) *Ignorability* means that we have measurements of all factors that materially concurrently affect the treatment decision (Day-3 vs. Day-5 transfers) and the outcome (implantation); *overlap* means that every embryo could have plausibly been transferred either on Day-3 or Day-5; SUTVA means that the failure or success of one embryo does not affect those of other embryos.

### Delineating the confounding contributions to the treatment effects of extended incubation to blastocyst transfer on embryo implantation

While evidence-based guidelines for determining embryo day-of-transfer are lacking, it is possible to outline the candidate factors that both direct the decision-making process regarding day-of-transfer and affect the implantation outcome. Most importantly, embryo quality directly affects implantation outcome and day-of-transfer decisions such that high quality embryos are favorably considered for extended incubation to blastocyst transfer over low-quality embryos, due to the associated risk of pre-blastulation developmental arrest. To decrease the risk of blastocyst-transfer cycle cancelation, the size of the reserve-set of embryos per oocyte retrieval that are valid candidates for transfer is another important parameter that may direct day-of-transfer decisions. This parameter is also associated with the implantation potential of the embryos that are selected for transfer and is thus expected to affect the implantation outcome.(*17*) In addition, maternal age is an important reproductive factor that is associated with an increase in chromosomal aberrations and is correlated with the size of the embryo reserve set per oocyte retrieval and with a decline in the developmental potential of implant.(*18, 19*) Hence, maternal age may also direct embryo transfer decision, including day-of-transfer.

We first consider the central embryonic and maternal factors that are expected to direct day-of-transfer decisions and affect implantation outcome, i.e. the potential confounders. To assess how embryo developmental potential correlates with day-of-transfer decisions, we employ the widely used morphokinetic profiling of the discrete transition events between distinctive developmental states as a metric of embryo quality.(*20-22*) The morphokinetic events (Fig. 2A-i) and the cell-cycle and synchronization intervals (Fig. 2A-ii) exhibited high degree of overlap between cleavage-stage and blastocyst transfers at 66 hours from fertilization. Similarly, there were negligible differences in the maternal age distributions between these transferred embryo sets (Fig. 2B). To assess the correlation between non-transferred reserve-set of embryos and day-of-transfer decisions, we separated between 8C^+^ embryos that consisted of ≥ 8 blastomers and 4C^-^ embryos that consisted of ≤ 4 blastomers at 66 hours from fertilization. In this manner, we addressed the potential confounding of high-quality embryos that were valid candidates for transfer and low-quality embryos. No statistically significant differences were found in the number of co-cultured 4C^-^ embryos (Fig. 2C-i). However, blastocyst transfers were enriched with a higher number of 8C^+^ embryo cycles (Fig. 2C-ii). Since the highest quality embryos are the first ones to be transferred, the lack of morphokinetic differences between cleavage-stage transferred embryos and transferred blastocyst indicates that most transfer cycles include at least one high quality embryo on Day-3. Furthermore, the fact that cleavage-stage and blastocyst transfers are not differentiated by maternal age implies that there are high quality embryos available for transfer that are generated both by younger and by older women. However, the number of high quality embryos is the parameter that appears to have a significant effect on day-of-transfer decisions and may also affect implantation outcome.(*23*) We thus conclude that we have measured or have proxies for most of the important and relevant confounders and proceed to discuss potential hidden confounders.

**Figure 2.**
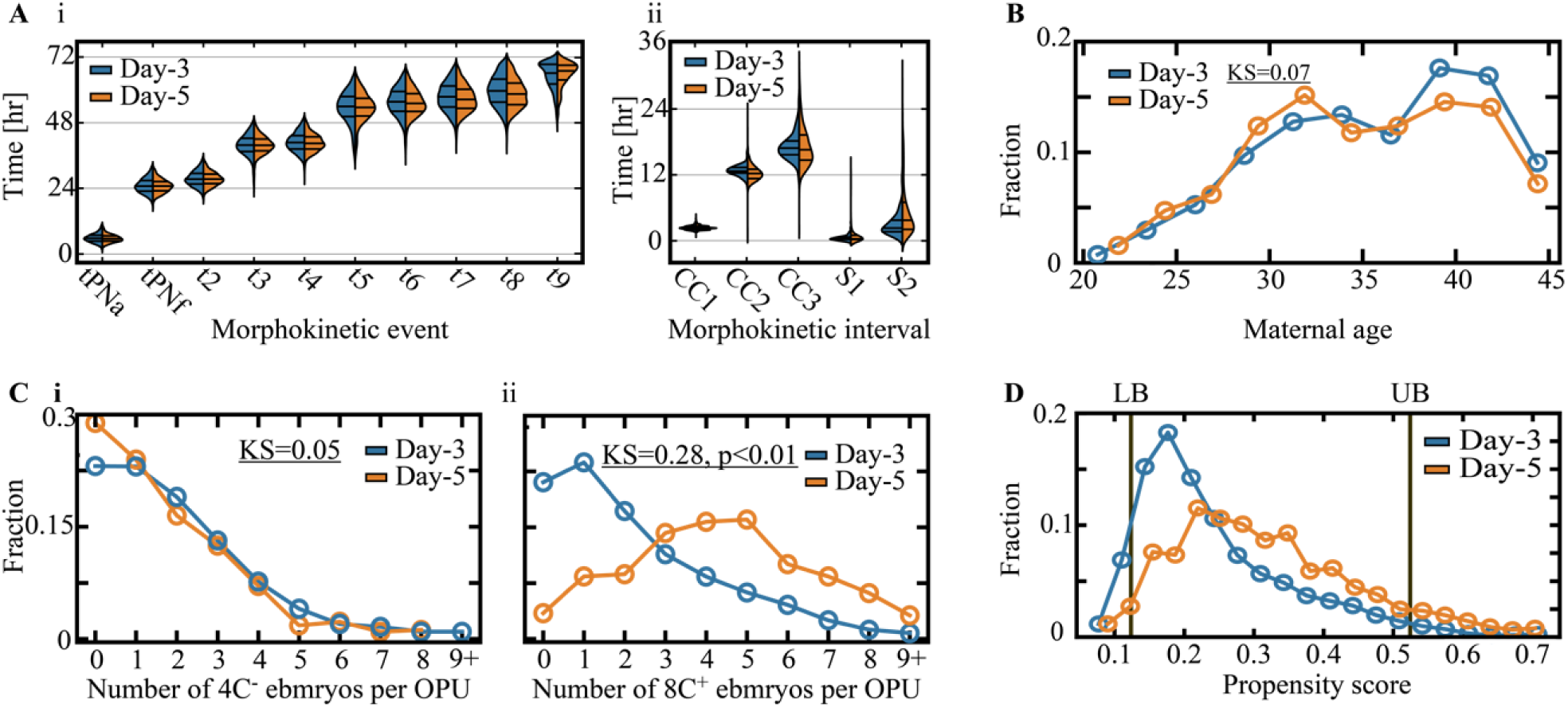
Feature analysis of cleavage-stage versus blastocyst transfers. **A**, No significant differences are observed between the temporal distributions of the (i) morphokinetic events and the (ii) cell cycle and synchronization intervals of Day-3 cleavage-stage transferred embryos (n=1,892) and Day-5 transferred blastocysts (n=799). KS distances < 0.1. Only high-quality embryos that reached 8-cell cleavage were included. **B**, The maternal age distributions of Day-3 and Day-5 transferred embryos are overlapping. **C**, The number of (i) low-quality (≤ 4 blastomeres) and (ii) high-quality (≥ 8 blastomeres) embryos of the same oocyte retrievals are compared between Day-3 and Day-5 freshly transferred cycles. Blastocyst transfers are associated with > 3 high-quality co-cultured embryos. **D**, Day-3 and Day-5 propensity score distributions were derived using a logistic regression prediction model of the day-of- transfer. Lower bound (LB) and upper bound (UB) values for excluding non-overlapping embryos are set by the 2.5 percentile and the 97.5 percentile of the Day-5 and Day-3 propensity score distributions. Abbreviations. ET: Embryo transfer. KS: Kolmogorov-Smirnov.

Counter to markers of embryo quality, maternal age, and oocyte retrieval reserve-set parameters, IVF history and clinical background factors are not included in the dataset and may thus contribute to hidden confounding. To validate that ignorability is satisfied, we consider the relationship between principal fertility-related conditions, day-of-transfer decision-making protocols, and embryo implantation outcome. It is expected that a history of implantation failures will bias the physician to switch the day-of-transfer whereas previous implantation successes would support transferring the embryo(s) at the same developmental stage. Notably, a history of negative implantation outcome was reported to correlate with a decline in future implantation potential.(*24*) Nevertheless, IVF history considerations are not included in the established guidelines and do not override the current recommendations for favoring single-blastocyst transfers under the defined range of conditions.(*25*) With respect to specific medical conditions, repeated implantation failure (RIF) is likely underlined by non-embryonic maternal aspects, including endometrial receptivity.(*26*) However, the potentially confounding effect is small with a reported 10% prevalence, which is likely over-diagnosed.(*27*) Unlike RIF, single blastocyst transfers are recommended for patients with past preterm delivery akin to the standard guidelines.(*28*) Similarly, single blastocyst transfers may be preferred in case of unicornuate uterus, which is a rare congenital condition that is characterized by an increased risk of miscarriage and preterm delivery, whereas no effects are known on the implantation potential in IVF treatments.(*29*) In summary, we surveyed the central indicators of IVF history and clinical background and demonstrated that their effects on day-of-transfer decisions and implantation outcome are either irrelevant or not significant. We therefore believe that all important factors affecting both treatment and outcome are represented in our dataset, leading us to conclude that the ignorability assumption is very nearly satisfied by our data-gathering process.

To explore the extent of *overlap* between cleavage-stage and blastocyst transfers, we fit a logistic regression model for predicting the day-of-transfer based on the parameters described above: (1) Embryo morphokinetic profiles. (2) Maternal age. (3) The number of co-cultured 4C^-^ embryos, and (4) 8C^+^ embryos per oocyte retrieval at 66 hours from fertilization. With respect to these feature vectors, cleavage-stage and blastocyst transfers were only partially separated. This is indicated by the propensity score, which quantifies the day-of-transfer prediction probability (Fig. 2D). We find that excluding (trimming(*30*)) the 10% of embryos with propensity below the 0.025 quantile of Day-5 transferred embryos or above 0.975 quantile of Day-3 transferred embryos removes all cases that lack counterexamples, thus satisfying the *overlap* assumption between conditions among the remaining embryos. There were no differences in the morphokientic profiles (Fig. 3A-i,ii), maternal age distributions (Fig. 3B-i), and number of 4C^-^ embryos per oocyte retrieval (Fig. 3B-ii) between the excluded and the remaining embryos. However, the excluded embryos are characterized either by a relatively small number (1) or very high number (9) of 8C^+^ embryos per oocytes retrieval (Fig. 3B-iii).

**Figure 3.**
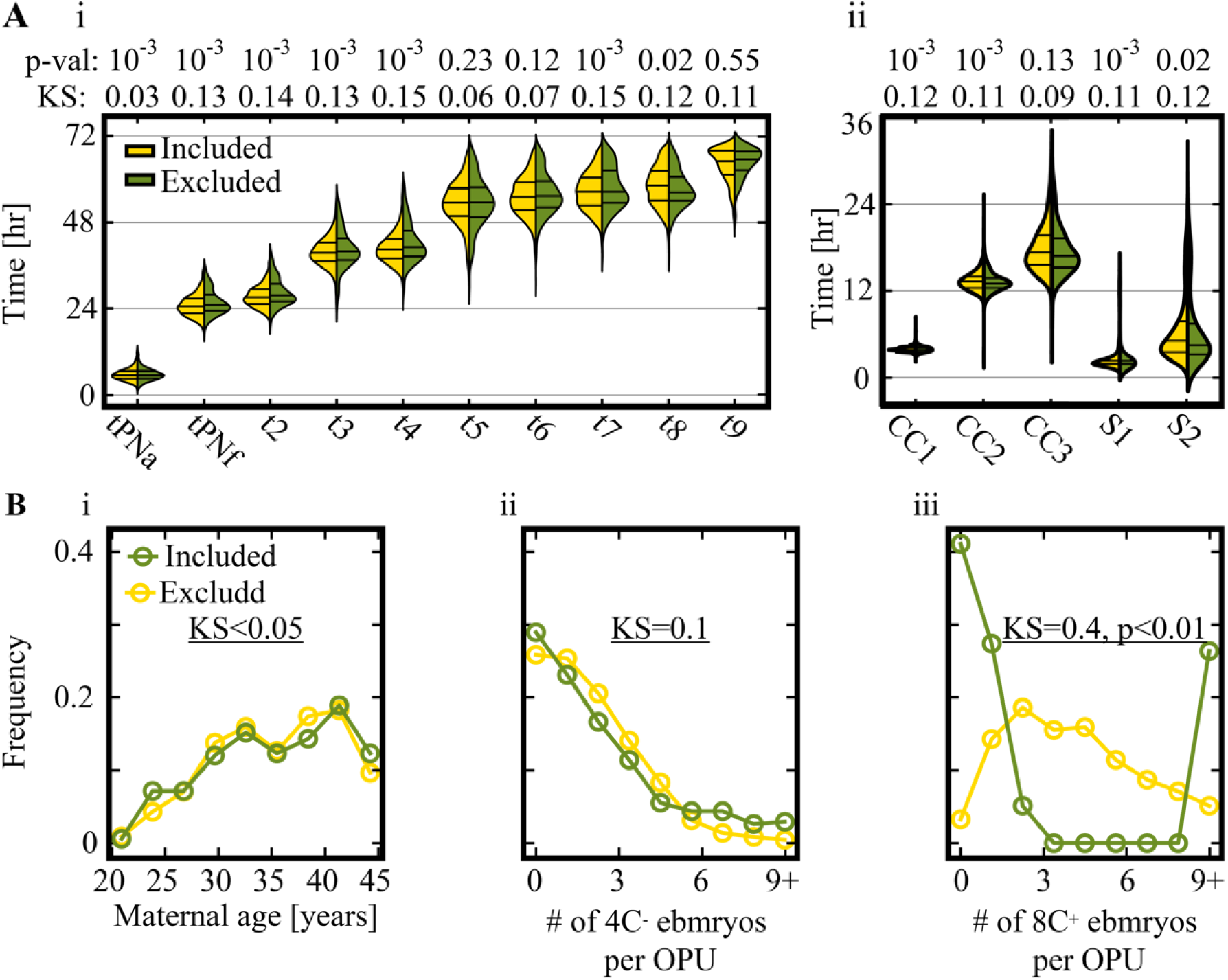
Statistical differences between embryos with propensity scores outside (excluded) and inside (included) the allowed propensity range. **(A)**, A comparison between the temporal distributions of the (i) morphokinetic events, and the (ii) cell cycle (CC1-to-CC3) and synchronization (S1, S2) intervals as evaluated 66 hours from fertilization of included versus excluded embryos. **(B)**, Comparison of (i) the maternal age distributions, (ii) number of low quality embryos (4C^-^ at 66 hours from fertilization) per oocyte retrieval, and (iii) number of high quality embryos (8C^+^ at 66 hours from fertilization) per oocyte retrieval of excluded versus included embryos. Included embryos: n=2,254 (Day-3) and n=829 (Day-5). Excluded embryos: n=251 (Day-3) and n=99 (Day-5). KS: Kolmogorov-Smirnov distance.

Similar to the ignorability and overlap assumptions, *SUTVA* is also satisfied by our data-gathering process since the outcome of the day-of-transfer treatment of one patient is independent of the transfer outcome of other patients. Hence, we have strong evidence that the dataset supports the possibility of valid causal inference for evaluating the causal effect of extended incubation and blastocyst transfer on implantation outcome. Our ability to infer these causal effects is also supported by the natural source of randomness related to day of OPU, as presented above.(*12, 31*)

### Evaluating the heterogeneous day-of-transfer treatment effect of extended incubation to blastocyst transfer on embryo implantation by fitting a causal forest model

Given the variation in the potential of embryos to blastulate, implant, and generate live birth, we hypothesized that the response to extended culture and to differences in endometrial synchronization that are associated with blastocyst transfers would also vary between embryos.(*32-34*) To evaluate the heterogeneous day-of-transfer treatment effect, we fit a causal forest (CF) model using the feature vectors of the embryos that we used for evaluating the propensity score. Using CF, we calculated the *Transfer Lift* of the embryos at 66 hours from fertilization, which is the estimated implantation potential (in terms of probability, between 0 and 1) of individual embryos if transferred on Day-5 minus the same potential if transferred on Day-3.(*35*) In this manner, the *Transfer Lift* provides a quantitative assessment of the heterogeneous day-of-transfer treatment effect. It is a property of individual embryos that is independent of whether the embryos are actually transferred or not. The *Transfer Lift* is bounded between -1 and 1, where positive *Transfer Lift* indicates that the implantation potential is predicted to be higher if transferred on Day-5 and vice versa. A mathematical description of the *Transfer Lift* is provided in the materials and methods section.

The mean *Transfer Lift* standard deviation of the test set and the train set embryos was 0.1 ± 0.07 and 0.1 ± 0.08, respectively, which quantifies the difference in the probability of implantation (treatment effect) between Day-5 and Day-3 transfers. While the majority of embryos obtained positive *Transfer Lift* with up to 0.3 higher estimated probability for implantation if transferred on Day-5, 6.3% of the test-set embryos and 8.5% of the train-set embryos obtained negative *Transfer Lift* with up to 0.1 higher estimated probability for implantation if transferred on Day-3 (Fig. 4A-i). Similar *Transfer Lift* distributions were obtained for embryos from different clinics (Fig. S1A-i,ii) and for single-versus-multiple embryo transfers (Fig. S1B-i,ii), thus demonstrating generality. As a control, we randomly permuted the implantation outcome (Fig. 4A-ii) and the day-of-transfer (Fig. 4A-iii), and fit CF models for each case. As expected, we obtained symmetric distributions with equal representation of positive and negative *Transfer Lift* embryos that were statistically-significantly separated from the non-permuted distributions. Importantly, there were no differences in the *Transfer Lift* distributions of cleavage-stage and blastocyst transferred embryos (Fig. 4B-i,ii).

**Figure 4.**
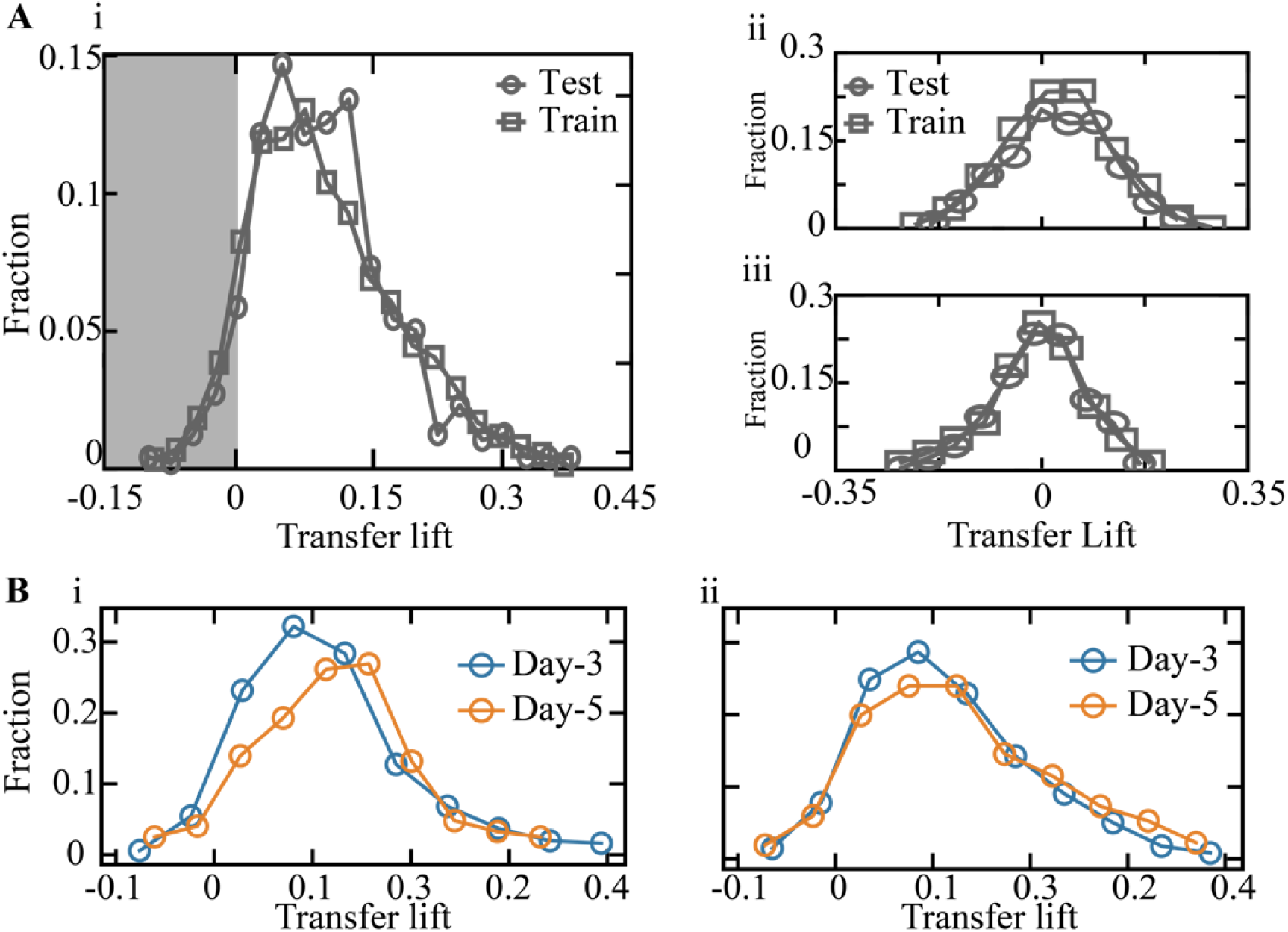
The *Transfer Lift* measures the difference in the implantation potential of embryos if transferred at blastocyst-stage relative to cleavage-stage. **A**, (i) The obtained test-set and train-set *Transfer Lift* distributions are asymmetric about zero, consisting of embryos with negative (gray background) and with positive (white background) *Transfer Lift* values. As a control, *Transferred Lift* was evaluated after randomly permuting (ii) the implantation outcome and (iii) the day-of-transfer labels. The differences of the implantation outcome (ii) and the day-of-transfer (iii) permuted distributions from the non-permuted Transfer Lift distribution (i), as evaluated using KS statistics, was 0.34 and 0.51 KS distance respectively, and p-value < 0.01. **B**, The *Transfer Lift* histogram of the (i) test set and (ii) train set embryos that were transferred on Day-3 and Day-5 overlap. Test set and train set: KS = 0.12. KS: Kolmogorov-Smirnov. STD: Standard deviation.

### Implementing embryo-specific day-of-transfer protocols is estimated to significantly improve implantation rate

Comparison of the morphokinetic profiles of 8C^+^ embryos reveals that negative *Transfer Lift* embryos were temporally-retarded relative to positive *Transfer Lift* embryos (Fig. 5A-i,ii). However, no distinctive maternal age dependence was detected (Fig. 5B-i,ii), no difference in the number of co-cultured 4C^-^ (Fig. 5C-i,ii) nor 8C^+^ (Fig. 5D-i,ii) embryos of the same oocyte retrieval reserve-set were found, and no statistically significant differences were observed in the Day-3 KIDScore values that are used for assessing implantation potential (Fig. 5E-i,ii).(*36*) Consistently, the adjusted mutual information between the *Transfer Lift* and the KIDScore values of test set and train set embryos was 0.005 and 0.013, respectively. These analyses indicate that the *Transfer Lift* is a novel property of individual embryos that is independent of maternal age, oocyte retrieval statistics, and predicted embryo quality.

**Figure 5.**
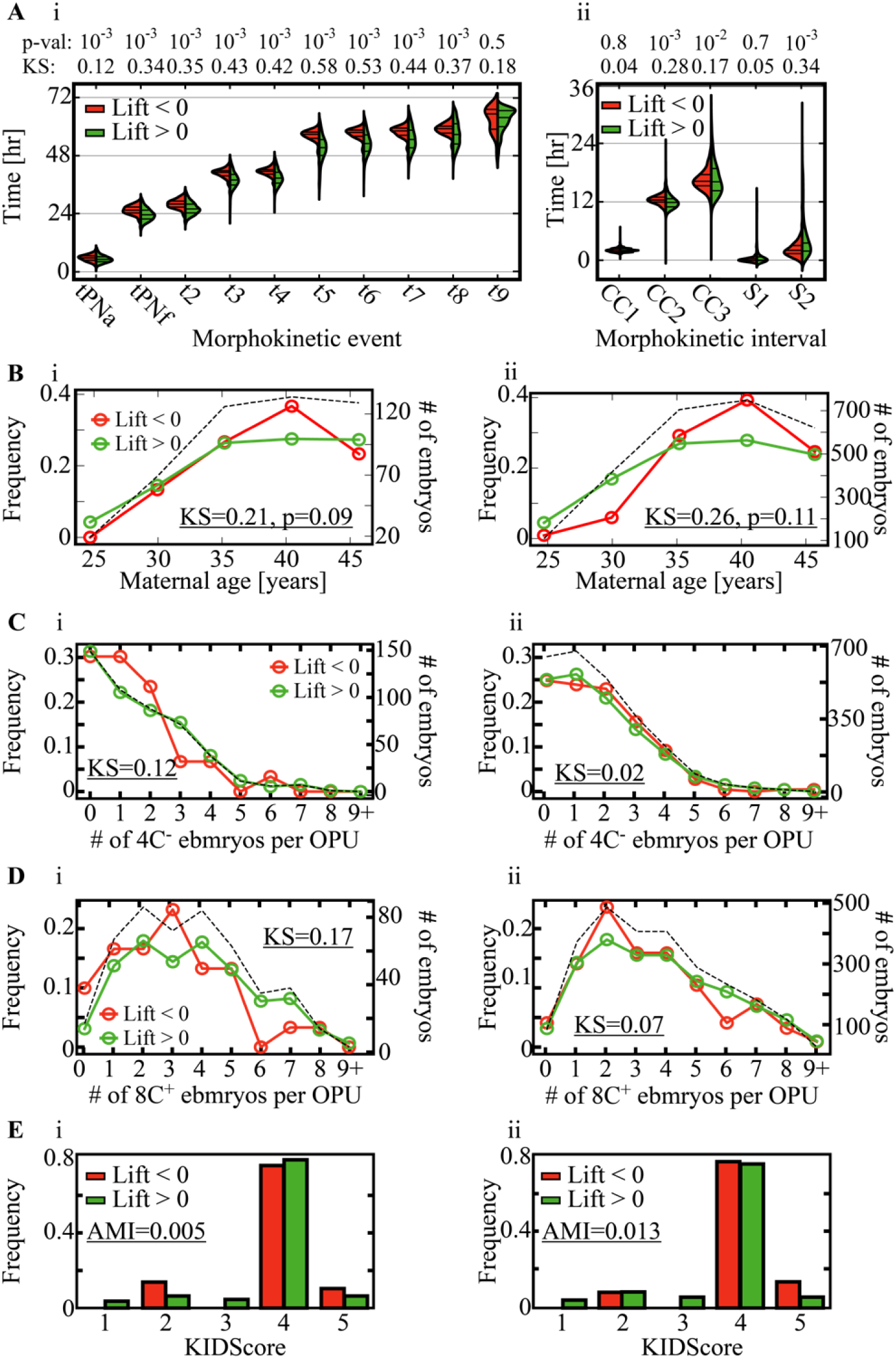
Feature analysis of positive versus negative *Transfer Lift* embryos. **A**, Comparison of the temporal distributions of the (i) morphokinetic events, and the (ii) cell cycles (CC1-to-CC3) and synchronization (S1, S2) intervals as evaluated 66 hours from fertilization demonstrate relative temporal retardation of the negative *Transfer Lift* embryos. **B**, Comparison of the maternal age distributions of (i) test-set and (ii) train-set embryos. **C**, Comparison of the number of low quality embryos (4C^-^ at 66 hours from fertilization) per oocyte retrieval of positive and negative *Transfer Lift* (i) test-set and (ii) train-set embryos. **D**, Comparison of the number of high quality embryos (8C^+^ at 66 hours from fertilization) per oocyte retrieval of positive and negative *Transfer Lift* (i) test-set and (ii) train-set embryos. **E**, Comparison of Day-3 KIDScore ranking of (i) test-set and (ii) train-set embryos. Dependence between the Transfer Lift distributions and the KIDScore distributions is quantified via AMI. The total number of embryos is depicted by the dashed lines in **B, C**, and **D**. Abbreviations. p-val: p-values. KS: Kolmogorov-Smirnov distances. AMI: Adjusted mutual information.

To explore the relationship between the *Transfer Lift* and the actual treatment outcome, we compared the implantation rates of negative and positive *Transfer Lift* test-set embryos. Satisfyingly, the average implantation rate of positive *Transfer Lift* embryos was higher when transferred on Day-5 and the average implantation rate of negative *Transfer Lift* embryos was higher when transferred on Day-3 (Fig. 6A-i). This was further validated for high-quality 8C^+^ embryos (Fig. 6A-ii). Our results thus strengthen the finding that the *Transfer Lift* measures the heterogeneous treatment effect and indicates that the implantation rate could be increased by transferring negative *Transfer Lift* embryos on Day-3 and positive *Transfer Lift* embryos on Day-5. To quantify the expected improvement in the implantation rate, we retrospectively re-adjusted the implantation outcome of positive *Transfer Lift* embryos that were transferred on Day-3 and negative *Transfer Lift* embryos that were transferred on Day-5 by adding their absolute *Transfer Lift* values. This re-adjustment scheme is based on the fact that the *Transfer Lift* measures the difference in probability of a given embryo to implant between cleavage-stage and blastocyst-stage transfer scenarios. Indeed, we estimate that transferring embryos according to their *Transfer Lift* would have increased the average implantation rate by 32% (Fig. 6B).

**Figure 6.**
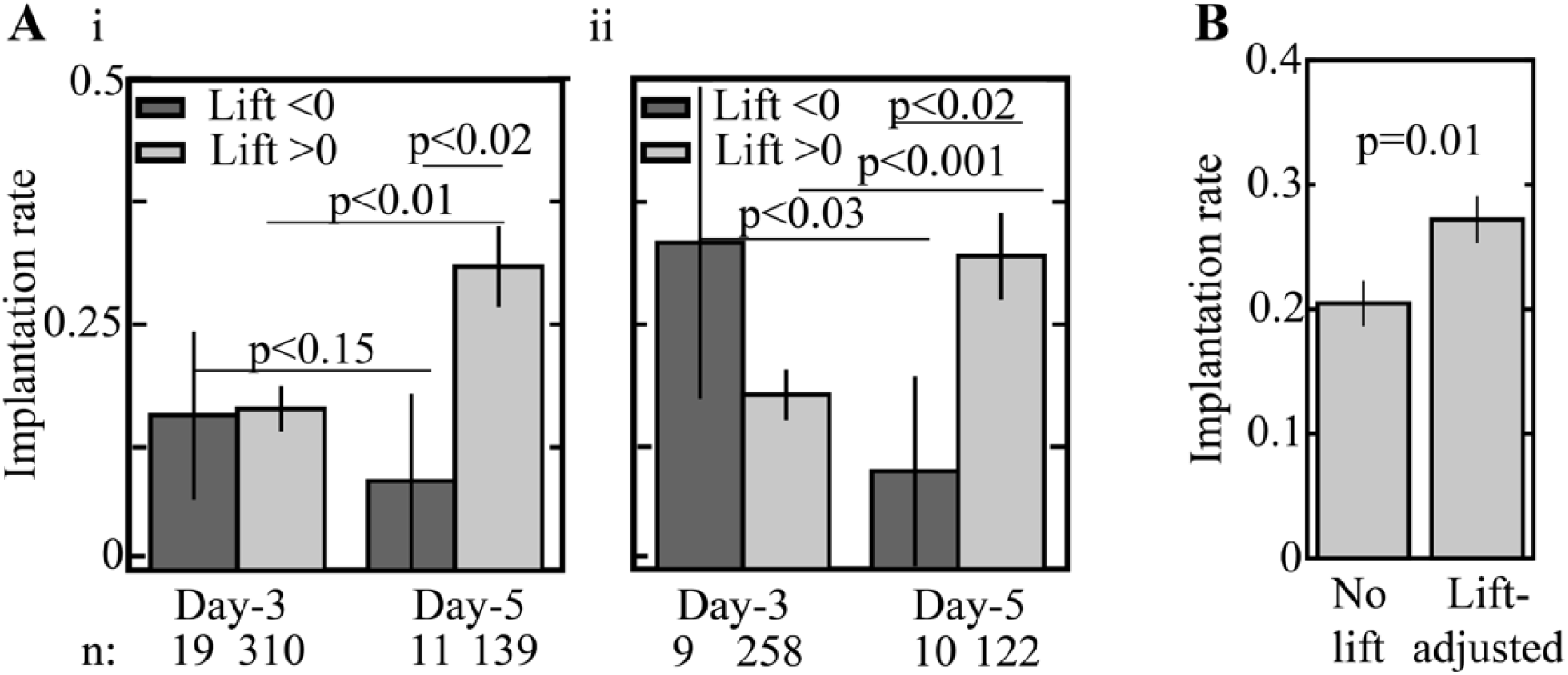
The *Transfer Lift* measures the difference in the implantation potential of embryos if transferred at blastocyst-stage relative to cleavage-stage. **A**, The implantation rates of negative and positive *Transfer Lift* embryos that were transferred on Day-3 and on Day-5 are compared. Average implantation rates are presented (i) for all test-set embryos, and (ii) across high-quality embryos only (8C+ at 66 hours). Error bars represent STD. **B**, The *Transfer Lift*-adjusted implantation rate accounts for the expected outcome of transferring negative and positive *Transfer Lift* embryos at cleavage-stage and blastocyst-stage, respectively. The average implantation outcome is estimated to improve by 32% relative to the current clinical practice (p-value 0.01). Error bars depict STD. KS: Kolmogorov-Smirnov. STD: Standard deviation.

## DISCUSSION

Incorporating evidence-based medicine principles in IVF-embryo transfer treatments is hindered by ethical constrains on clinical trials and bias in observational studies.(*37*) Using an expansive retrospective multicenter dataset, we confirm the current clinical practice by which extended culture to blastocyst transfer is considered in case the size of the reserve-set of high-quality embryos per oocyte retrieval is high, thus decreasing the risk that no blastocysts would be generated by Day-5 leading to transfer cycle cancelation. However, the response of the embryos that are selected for transfer to extended culture for blastocyst transfer is the determinant factor. Hence, we considered the potential confounders and hidden confounders, identified a natural source of randomness, and demonstrated the likelihood that the required causal assumptions are satisfied by the data-gathering process and can thus support applying causal inference methods to calculate the heterogeneous day-of-transfer treatment effect on implantation potential.(*12, 38, 39*) By fitting a causal forest model, we evaluated the *Transfer Lift*, which measures the heterogeneous treatment effect of extended culture to blastocyst transfer on the implantation outcome. Akin to the developmental potential to implant and generate live birth, the *Transfer Lift* is a quantitative property of the embryos that accounts for the difference in the probability to implantation between blastocyst and cleavage-stage transfer scenarios.

The molecular mechanisms that underlie the *Transfer Lift* cannot be determined via a computational vision-based approach and are thus beyond the scope of this work. However, differences in the *Transfer Lift* suggest that embryos respond heterogeneously to the environmental conditions that are set by the culture medium and/or in the uterus. Specifically, negative *Transfer Lift* embryos are found to be morphokinetically-retarded relative to positive *Transfer Lift*, which might introduce differences in embryo-endometrium synchrony. Compared with positive *Transfer Lift* embryos, negative *Transfer Lift* embryos are likely more sensitive to the extended culture environment during morula compaction and blastulation. On the other hand, positive *Transfer Lift* embryos might gain from a better synchronization with the uterine environment if transferred as blastocysts during the window of implantation.(*32*) In terms of medical utility, we present a decision support tool for estimating the patient-level day-of-transfer treatment effect in a manner that is complementary with existing classifiers of embryo implantation potential(*13*) and 1^st^ trimester miscarriage potential.(*40*) We estimate that optimizing the day-of-transfer of the embryos that are selected for transfer could significantly improve implantation rate, thus supporting single embryo transfer policy while shortening time-to-pregnancy. Finally, our work relies on observational data; despite our best efforts and the uniqueness of our dataset, we can never rule out the possibility of some unmeasured confounder inducing spurious correlations misleading our model. We hope our work will spur clinical experiments based on per-embryo treatment protocols.

## MATERIALS AND METHODS

### Dataset

In this work we used morphokinetic profiles of preimplantation embryos and related clinical labels as previously reported.(*13*) In short, the morphokientic profiles and the clinical labels were obtained using the dataset that we assembled and consisted of time-lapse imaging video files of 2,175 fresh cycles and approved by the Investigation Review Boards of the data-providing medical centers: Hadassah Hebrew University Medical center IRB number HMO 558-14; Kaplan Medical Center IRB 0040-16-KMC; Soroka Medical Center IRB 0328-17-SOR; Rabin Medical Center IRB 0767-15-RMC. Here we included data from 3,433 ICSI-fertilized embryos that were transferred to the uterus either on Day-3 or on Day-5 from time of fertilization with known implantation outcome supplemented with 12,907 non-transferred co-cultured embryos (Table-1). Clinical metadata included maternal age labels. The embryos were cultured in twelve time-lapse incubations systems (Embryoscope, Vitrolife) in our medical centers. The transferred embryos were randomly divided into train (85%) and test (15%) sets.

**Table-1:**
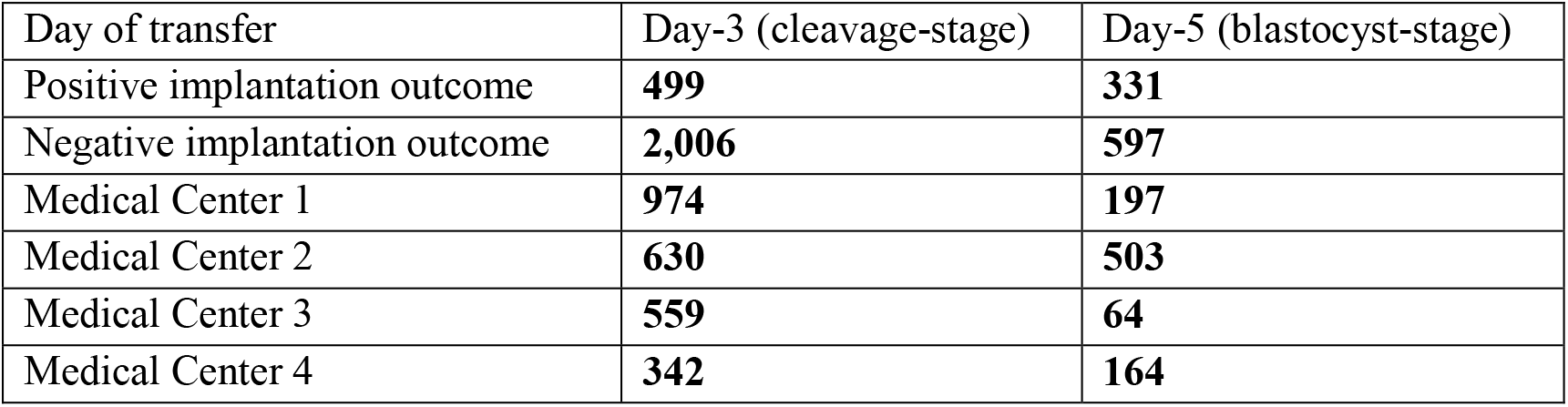
Cleavage-stage and blastocyst-stage transferred embryos.

### Computational modeling

Propensity score was calculated using a logistic regression model (scikit-learn python package), with L2 penalty. A CF model was developed using the Generalized Random Forests package(*35*) that is implemented in the R programming language.(*41*) The model parameters were selected based on a fivefold cross-validation scheme. CF training was performed using 300 trees and a minimum of 20 embryos per leaf.

### Fitting a causal forest model

For a given embryo *i*, the feature vector *X*_*i*_ consists of the morphokientic profile, maternal age, and the number of high and low quality co-cultured embryo at 66 hours from fertilization as defined in the main text (Fig. S2A). The treatment indicator divides the embryos into “non-treated” Day-3 transferred embryos (*W*_*i*_ *= 0*) and “treated” Day-5 transferred embryos (*W*_*i*_ *= 1*). The individual treatment response *Y*_*i*_ corresponds to negative implantation (*Y*_*i*_ = 0) or positive implantation (*Y*_*i*_= 1) outcome. Using feature vectors *X*_*i*_, we trained a CF model consisting of *B* causal decision trees (CDTs; Fig. S2B).(*35*) Co-representation of Day-3 and Day-5 transferred embryos was verified along the entire feature vector range by excluding 10% of the embryos that lacked counterexamples with overlapping propensity scores (Fig. 2D). Heuristically, each CDT divides the embryos according to *X*_*i*_ with respect to *W*_*i*_ into discrete leaves (Fig. S2C). Within a given leaf 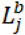, the treatment effect 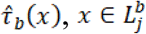, is defined by the average response of the treated embryos minus the average response of the non-treated embryos:

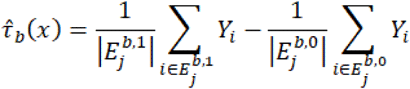

where 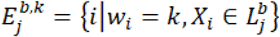. The conditional average treatment effect of any embryo with features is evaluated by the *Transfer Lift*, which is the averaged treatment effect across all *B* causal trees in the forest:

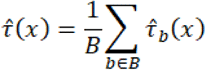

The *Transfer Lift* measures the difference in the implantation potential in case it was transferred on Day-5 relative to Day-3. The *Transfer Lift* ranges between -1 and 1, where positive values account for embryos whose implantation potential is higher if transferred on Day-5 and vice versa.

### Statistical analysis

Graphs and heatmaps were plotted using matplotlib and seaborn python packages. Student’s t-test, Wilcoxon rank-sum test, and Kolmogorov-Smirnov statistics were calculated using the Python SciPy package. Adjusted mutual information was calculated using the Scikit-learn python package with the arithmetic average method. The statistical significance of the differences in implantation rates between Day-3 and Day-5 transferred embryos with negative and positive *Transfer Lift* (Fig. 6A) was determined using a permutation test. The null hypothesis was that the observed, low implantation rate in the smaller group is due to chance. In each test, 1,000 permutations were randomly sampled. Our results are validated using a Wilcoxon rank-sum test that generated comparable results.

## Acknowledgments

We are thankful to Prof. Liran Carmel for fruitful discussion and to our original data providers from Kaplan Medical Center, Dr. Yuval Or and Prof. Zeev Shoham; Hadassah Medical Center, Dr. Assaf Ben-Meir and Dr. Arye Hurwit; Soroka University Medical Center, Dr. Iris Har-Vardi; and Rabin Medical Center, Dr. Yoel Shufaro, as credited by Kan-Tor et al.

## Author contributions

Conceptualization: YKT and AB

Methodology: YKT, MG, US, and AB.

Investigation: YKT, NS, and AB.

Visualization: YKT.

Funding acquisition: AB.

Project administration: AB.

Supervision: AB.

Writing – original draft: YKT, US and AB.

Writing – review & editing: NA.

## Data and materials availability

Morphokinetic profiles and code used in the analysis will be made available on GitHub for purposes of reproducing or extending the analysis.

## SUPPLEMENTARY MATERIAL

**Figure S1.**
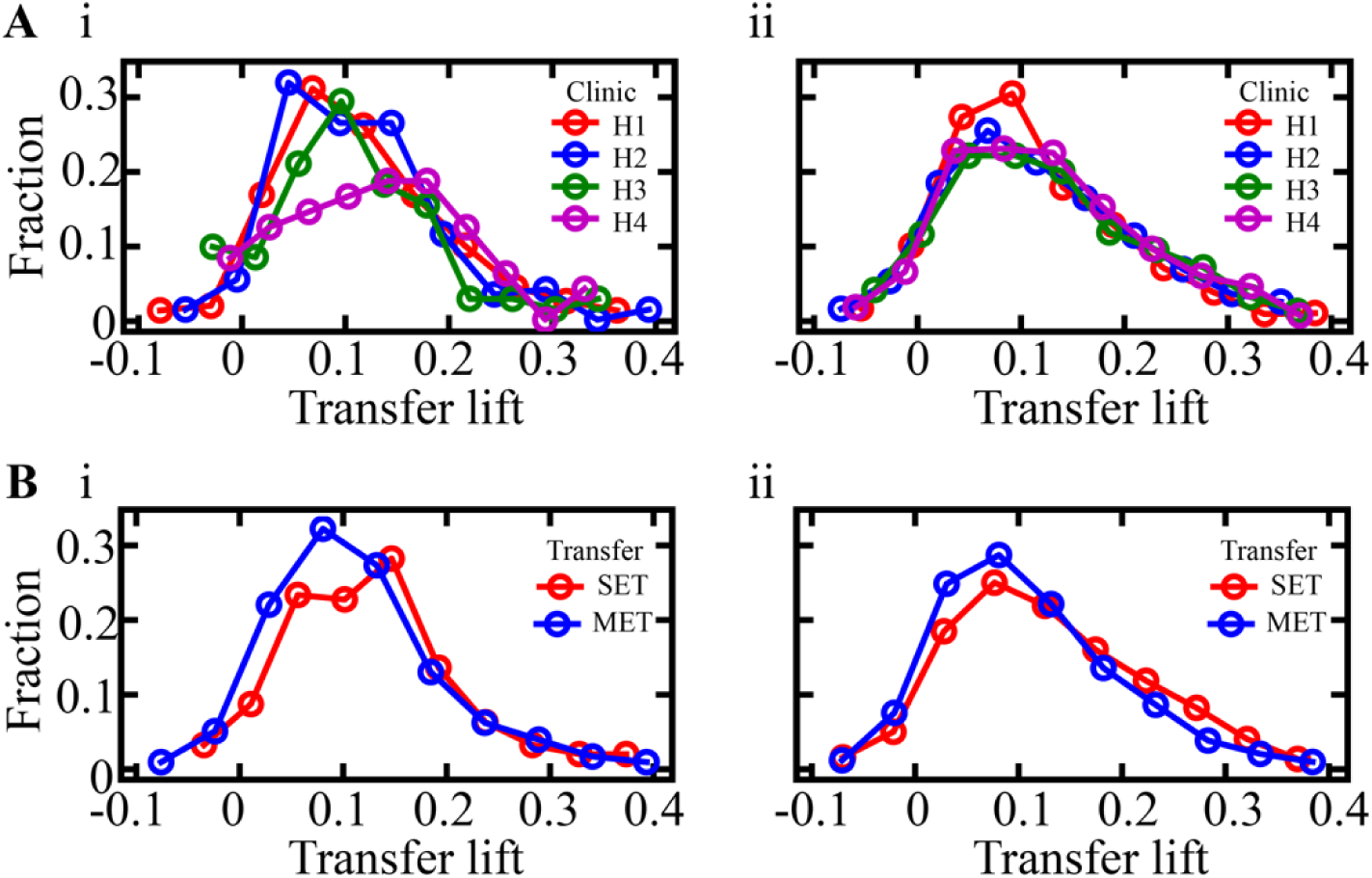
The *Transfer Lift* is stable across medical and clinical parameters: **A** Comparison of the *Transfer Lift* distributions of the embryos that were obtained from H1-through-H4 IVF clinics. Peak-to-peak KS distance across clinics of (i) test-set and (ii) train set embryos was 0.24 and 0.12, respectively. **B**, Comparison of the *Transfer Lift* distributions of single-embryo transfers (SETs) and multiple-embryo transfers (METs). KS distance between SET and MET of (i) test set and (ii) train set embryos was 0.17 and 0.11, respectively. KS: Kolmogorov-Smirnov.

**Figure S2.**
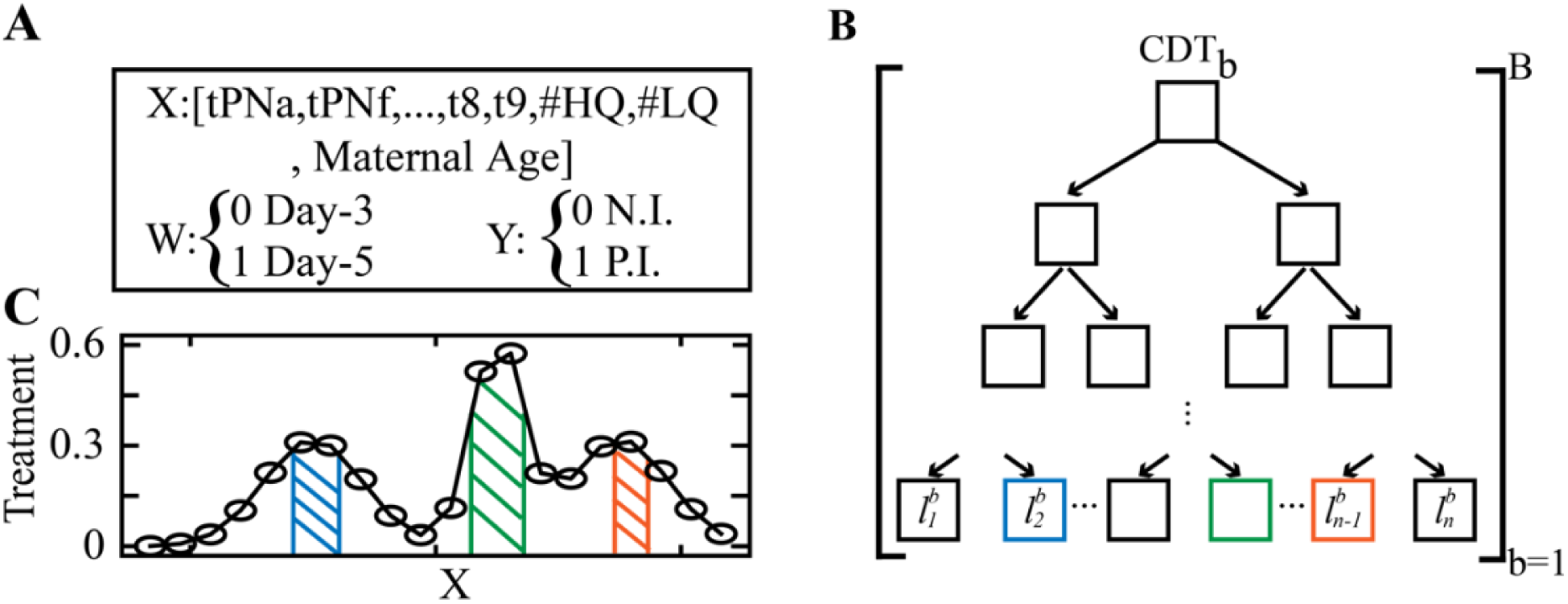
Illustration of the causal forest model: **A**, The counterfactual inference problem is set by the embryo feature vector *X*, the treatment indicator *W*, and the treatment response *Y*. **B**, Schematic illustration of a single causal decision tree (CDT), among a total of *B* CDTs, illustrates the hierarchical classification of the embryos into discrete leaves. **C**, Heuristic representation depicts the variation in the Treatment *W* between clusters of embryos (leaves) of similar feature profiles *X*.

## REFERENCES

1. N. Gleicher, V. A. Kushnir, D. H. Barad, Worldwide decline of IVF birth rates and its probable causes. Hum Reprod Open 2019, hoz017 (2019).

2. N. Gleicher, V. A. Kushnir, D. F. Albertini, D. H. Barad, Improvements in IVF in women of advanced age. J Endocrinol 230, F1–6 (2016).

3. D. K. Gardner, M. Lane, Culture of viable human blastocysts in defined sequential serum-free media. Hum Reprod 13 Suppl 3, 148-159; discussion 160 (1998).

4. D. K. Gardner et al., Single blastocyst transfer: a prospective randomized trial. Fertil Steril 81, 551–555 (2004).

5. A. Criniti et al., Elective single blastocyst transfer reduces twin rates without compromising pregnancy rates. Fertil Steril 84, 1613–1619 (2005).

6. D. M. Kissin et al., Embryo transfer practices and multiple births resulting from assisted reproductive technology: an opportunity for prevention. Fertil Steril 103, 954–961 (2015).

7. D. Glujovsky, C. Farquhar, Cleavage-stage or blastocyst transfer: what are the benefits and harms? Fertil Steril 106, 244–250 (2016).

8. D. Glujovsky, C. Farquhar, A. M. Quinteiro Retamar, C. R. Alvarez Sedo, D. Blake, Cleavage stage versus blastocyst stage embryo transfer in assisted reproductive technology. Cochrane Database Syst Rev, CD002118 (2016).

9. J. Wilkinson et al., Don’t abandon RCTs in IVF. We don’t even understand them. Human Reproduction 34, 2093–2098 (2019).

10. A. Arav, A. Aroyo, S. Yavin, Z. Roth, Prediction of embryonic developmental competence by time-lapse observation and ‘shortest-half’ analysis. Reprod Biomed Online 17, 669–675 (2008).

11. J. G. Lemmen, I. Agerholm, S. Ziebe, Kinetic markers of human embryo quality using time-lapse recordings of IVF/ICSI-fertilized oocytes. Reprod Biomed Online 17, 385–391 (2008).

12. R. J. Hernán MA, Causal Inference: What If. (Boca Raton: Chapman & Hall/CRC).

13. Y. Kan-Tor et al., Automated Evaluation of Human Embryo Blastulation and Implantation Potential using Deep-Learning. Advanced Intelligent Systems, 202000080 (2020).

14. A. Abbara et al., Follicle Size on Day of Trigger Most Likely to Yield a Mature Oocyte. Front Endocrinol (Lausanne) 9, 193 (2018).

15. R. B. D’Agostino, Jr., R. B. D’Agostino, Sr., Estimating treatment effects using observational data. JAMA 297, 314–316 (2007).

16. D. B. Rubin, Causal inference using potential outcomes: Design, modeling, decisions. J Am Stat Assoc 100, 322–331 (2005).

17. C. Racowsky et al., The number of eight-cell embryos is a key determinant for selecting day 3 or day 5 transfer. Fertil Steril 73, 558–564 (2000).

18. J. R. Gruhn et al., Chromosome errors in human eggs shape natural fertility over reproductive life span. Science 365, 1466–1469 (2019).

19. S. Munne et al., Maternal age, morphology, development and chromosome abnormalities in over 6000 cleavage-stage embryos. Reprod Biomed Online 14, 628–634 (2007).

20. D. K. Gardner, M. Meseguer, C. Rubio, N. R. Treff, Diagnosis of human preimplantation embryo viability. Hum Reprod Update 21, 727–747 (2015).

21. G. Coticchio et al., Embryo morphokinetic score is associated with biomarkers of developmental competence and implantation. J Assist Reprod Genet, (2021).

22. Y. Motato et al., Morphokinetic analysis and embryonic prediction for blastocyst formation through an integrated time-lapse system. Fertil Steril 105, 376–384 e379 (2016).

23. F. Devreker et al., Selection of good embryos for transfer depends on embryo cohort size: implications for the ‘mild ovarian stimulation’ debate. Hum Reprod 14, 3002–3008 (1999).

24. A. Smith, K. Tilling, S. M. Nelson, D. A. Lawlor, Live-Birth Rate Associated With Repeat In Vitro Fertilization Treatment Cycles. JAMA 314, 2654–2662 (2015).

25. M. Practice Committee of the American Society for Reproductive, a. a. o. Practice Committee of the Society for Assisted Reproductive Technology. Electronic address, Blastocyst culture and transfer in clinically assisted reproduction: a committee opinion. Fertil Steril 110, 1246–1252 (2018).

26. R. J. Paulson, M. V. Sauer, R. A. Lobo, Factors affecting embryo implantation after human in vitro fertilization: a hypothesis. Am J Obstet Gynecol 163, 2020–2023 (1990).

27. E. Somigliana et al., Repeated implantation failure at the crossroad between statistics, clinics and over-diagnosis. Reprod Biomed Online 36, 32–38 (2018).

28. A. a. o. Practice Committee of the American Society for Reproductive Medicine. Electronic address, T. Practice Committee of the Society for Assisted Reproductive, Guidance on the limits to the number of embryos to transfer: a committee opinion. Fertil Steril 107, 901–903 (2017).

29. Y. Chen, V. Nisenblat, P. Yang, X. Zhang, C. Ma, Reproductive outcomes in women with unicornuate uterus undergoing in vitro fertilization: a nested case-control retrospective study. Reprod Biol Endocrinol 16, 64 (2018).

30. M. Caliendo, S. Kopeinig, Some practical guidance for the implementation of propensity score matching. J Econ Surv 22, 31–72 (2008).

31. J. Pearl, An Introduction to Causal Inference. Int J Biostat 6, (2010).

32. M. Ruiz-Alonso, D. Valbuena, C. Gomez, J. Cuzzi, C. Simon, Endometrial Receptivity Analysis (ERA): data versus opinions. Hum Reprod Open 2021, hoab011 (2021).

33. B. A. Market-Velker, A. D. Fernandes, M. R. Mann, Side-by-side comparison of five commercial media systems in a mouse model: suboptimal in vitro culture interferes with imprint maintenance. Biol Reprod 83, 938–950 (2010).

34. R. J. Chason, J. Csokmay, J. H. Segars, A. H. DeCherney, D. R. Armant, Environmental and epigenetic effects upon preimplantation embryo metabolism and development. Trends Endocrinol Metab 22, 412–420 (2011).

35. S. Athey, J. Tibshirani, S. Wager, Generalized Random Forests. Ann Stat 47, 1148–1178 (2019).

36. B. M. Petersen, M. Boel, M. Montag, D. K. Gardner, Development of a generally applicable morphokinetic algorithm capable of predicting the implantation potential of embryos transferred on Day 3. Hum Reprod 31, 2231–2244 (2016).

37. N. Gleicher, D. H. Barad, Misplaced obsession with prospectively randomized studies. Reprod Biomed Online 21, 440–443 (2010).

38. U. Shalit, Can we learn individual-level treatment policies from clinical data? Biostatistics 21, 359–362 (2020).

39. J. Pearl, Causal inference in statistics: An overview. Statistics Surveys 3, 96-146, 151 (2009).

40. T. Amitai, Y. Kan-Tor, N. Srebnik, A. Buxboim, Embryo classification beyond pregnancy: Early prediction of first trimester miscarriage using machine learning. medRxiv, (2020).

41. R. D. C. Team. (R Foundation for Statistical Computing, Vienna Austria, 2010).

